# Phosphorylation of aldose-6-phosphate reductase from *Prunus persica* leaves

**DOI:** 10.1101/2022.07.05.498475

**Authors:** Matías D. Hartman, Bruno E. Rojas, Danisa M. L. Ferrero, Alejandro Leyva, Rosario Durán, Alberto A. Iglesias, Carlos M. Figueroa

## Abstract

Sugar-alcohols are major photosynthates in plants from the Rosaceae family. Expression of the gene encoding aldose-6-phosphate reductase (Ald6PRase), the critical enzyme for glucitol synthesis in rosaceous species, is regulated by physiological and environmental cues. Additionally, Ald6PRase is inhibited by small molecules (hexose-phosphates and inorganic orthophosphate) and oxidizing compounds. This work demonstrates that Ald6PRase from peach leaves is phosphorylated *in planta* at the N-terminus. We also show *in vitro* phosphorylation of recombinant Ald6PRase by a partially purified kinase extract from peach leaves containing Ca^2+^-dependent protein kinases (CDPKs). Moreover, phosphorylation of recombinant Ald6PRase was inhibited by hexose-phosphates, phosphoenolpyruvate and pyrophosphate. We further show that phosphorylation of recombinant Ald6PRase was maximal using recombinant CDPKs. Overall, our results suggest that phosphorylation could fine-tune the activity of Ald6PRase.

## 1. Introduction

In addition to sucrose and starch, sugar-alcohols are major photosynthetic products in plants from the Rosaceae family, like apple, pear, and peach, among others (Lewis and Smith, 1967). Glucitol (Gol), also known as sorbitol, is a sugar-alcohol produced in mature leaves from glucose 6-phosphate (Glc6P) after the combined action of aldose-6-phosphate reductase (Ald6PRase; EC 1.1.1.200), a member of the aldo-keto reductase (AKR) superfamily, and Gol6P phosphatase (EC 3.1.3.50). Gol is then translocated to heterotrophic tissues, where it is oxidized to fructose (Fru) by Gol dehydrogenase (GolDHase; EC 1.1.1.14; Figueroa et al., 2016; Loescher and Everard, 2004). Research performed in the last two decades established that Gol metabolism is regulated at the post-transcriptional level (Kanayama et al., 2006; Lloret et al., 2017; Lou et al., 2018; Suzuki and Dandekar, 2014). Additionally, we demonstrated that redox regulation of peach Ald6PRase and GolDHase orchestrates Gol metabolism in both source and sink tissues, respectively (Hartman et al., 2017, 2014).

More than 450 distinct protein post-translational modifications (PTMs) have been described so far (Khoury et al., 2011) and it is increasingly evident that many proteins, perhaps all, have multiple PTMs regulating their biological activity, subcellular location, and interaction with other proteins and/or nucleic acids (Hunter, 2007). Many plant enzymes are regulated by phosphorylation (Duncan et al., 2006; Gregory et al., 2009; McMichael et al., 1993; Piattoni et al., 2011); however, only a few works have dealt with protein phosphorylation in rosaceous species (Wang et al., 2014; Yu et al., 2021). Interestingly, a recent report (Yu et al., 2021) showed that phosphorylation of GolDHase by SNF1-related kinase 1 (SnRK1) promotes Gol metabolism and accumulation of sugars in peach fruits.

In this work, we demonstrate that Ald6PRase from peach leaves is phosphorylated *in planta* at the N-terminus. Additionally, we show that a partially purified kinase extract (PPKE) from peach leaves *in vitro* phosphorylates recombinant peach Ald6PRase (*Ppe*Ald6PRase). Finally, we prove that *Ppe*Ald6PRase is preferentially phosphorylated by Ca^2+^-dependent protein kinases (CDPKs). Overall, our work suggets that phosphorylation of Ald6PRase could be an important mechanism for regulating sugar-alcohol synthesis in rosaceous species.

## 2. Materials and methods

### 2.1 Plant material, bacterial strains, and reagents

Mature leaves from peach (*Prunus persica* cv. Flordaking) trees were harvested between 8 and 10 am at Campo Experimental en Cultivos Intensivos y Forestales (Facultad de Ciencias Agrarias, Universidad Nacional del Litoral, Santa Fe, Argentina), immediately frozen with liquid N_2_ and stored at -80°C until use. *Escherichia coli* TOP10 cells (Invitrogen) were used for cloning procedures and plasmid maintenance. Protein expression was carried out using *E. coli* BL21 Star (DE3) cells (Invitrogen). The substrates employed to measure enzymatic activities were from Sigma Aldrich. All other chemicals were of the highest quality available.

### 2.2 Protein extraction

Plant material was homogenized to a fine powder in liquid nitrogen with a mortar and pestle. For native protein extraction, 20 mg of fresh weight (FW) powdered tissue were extracted with 200 µl of a buffer containing 50 mM MOPS-NaOH pH 7.5, 0.1% (v/v) Triton X-100, 20% (v/v) glycerol, 4% (w/v) polyethylene glycol 8000 (PEG8000), 1 mM dithiothreitol (DTT), 5 mM MgCl_2_, 1 mM EDTA, 1% (w/v) polyvinylpolypyrrolidone, and Halt Protease and Phosphatase Inhibitor Cocktail (Thermo Fisher Scientific). In the case of denatured protein extraction, 20 mg of powdered FW tissue were extracted with 200 µl of the previously described buffer supplemented with 4.5 M urea and 4% (v/v) Triton X-100. In both cases, samples were treated for 30 min at 4°C with gentle homogenization and then centrifuged at 15,000 x *g* for 15 min at 4°C. The resulting supernatants were separated from cellular debris and used immediately. Protein content was determined using the Bradford reagent (Bradford, 1976).

### 2.3 Dephosphorylation assays

Proteins were incubated with 100 mM Tris-HCl pH 8.5, 1 mM EDTA, 10 mM MgCl_2_, 1.2 mM CaCl_2_, 20 mM 2-mercaptoethanol and 1 mM phenylmethylsulfonyl fluoride (PMSF); or with the same buffer supplemented with 5 U of calf intestine alkaline phosphatase (CIAP) from Promega. Samples were incubated at 30°C for 1 h. Reactions were stopped by the addition of SDS-PAGE sample buffer.

### 2.4 Enrichment of phosphoproteins

Enrichment of phosphorylated proteins was done as previously described by (Muszyńska et al., 1986). Briefly, 1 g of powdered leaf tissue was extracted with an ice-cold buffer containing 50 mM Hepes-KOH pH 7.5, 10% (v/v) glycerol, 0.25% (w/v) BSA, 0.1% (v/v) Triton X-100, 10 mM MgCl_2_, 1 mM EDTA, 1 mM EGTA, 1 mM PMSF and 1 mM DTT. The mixture was incubated on ice for 20 min with constant homogenization. The resulting extract was centrifuged for 30 min at 4°C and 15,000 x *g*. The supernatant was recovered and an aliquot containing 400 µg of total proteins was supplemented with 1.2 ml of 50 mM MES-NaOH pH 6.0. The resulting sample was incubated with 100 µl iminodiacetic acid-Fe^3+^ (IMAC-Fe^3+^) previously equilibrated with 50 mM MES-NaOH pH 6.0 for 1 h at 4°C with constant agitation. Non-adsorbed proteins were removed by washing twice with 2 ml of 50 mM MES-NaOH pH 6.0. The phosphorylated proteins were eluted in three steps by increasing the pH. Elutions were performed as follows: (1) five-column volumes of 50 mM PIPES-HCl pH 7.2; (2) three-column volumes of 50 mM Tris-HCl pH 8.0; and (3) two-column volumes of 50 mM Tris-HCl pH 9.0. Proteins eluted at pH 9.0 were further treated with 5 U of CIAP (Promega) and the buffer provided by the manufacturer (a negative control was prepared without phosphatase). The resulting samples were incubated at 30°C for 1 h and reactions were stopped by adding SDS-PAGE sample buffer.

### 2.5 Partial purification of protein kinases

Protein kinases from peach leaves were partially purified as previously described (Piattoni et al., 2011; Toroser et al., 2000). All purification procedures were performed at 4°C. Briefly, 25 g FW of leaves were extracted with 100 ml of a buffer containing 50 mM MOPS-NaOH pH 7.5, 2 mM EGTA, 2 mM EDTA, 5 mM DTT, 0.5 mM PMSF, 25 mM NaF, and 0.1% (v/v) Triton X-100. The sample was centrifuged at 10,000 x *g* for 15 min; then, PEG8000 was added to the supernatant to a final concentration of 3% (w/v). The resulting suspension was stirred for 10 min and then centrifuged at 10,000 x *g* for 10 min. Afterward, the supernatant was adjusted to 20% (w/v) PEG8000, stirred for 15 min, and centrifuged as before. The precipitate was resuspended in a buffer containing 12.5 ml of 50 mM MOPS-NaOH pH 7.5, 2 mM EGTA, 2 mM EDTA, 5 mM Na_4_P_2_O_7_, 5 mM NaF, and 2.5 mM DTT. The suspension was clarified by centrifugation at 10,000 x *g* for 10 min, and the supernatant was loaded onto a 2-ml Q-Sepharose column (Amersham Pharmacia Biotech). The column was washed with 20 column-volumes of Buffer A (50 mM MOPS-NaOH pH 7.5 and 1 mM DTT). Proteins were eluted with a 70-ml linear gradient of NaCl in Buffer A (from 0 to 500 mM). Collected fractions were assayed for protein kinase activity as described in section 2.6, using *Ppe*Ald6PRase as substrate. Active fractions were pooled and supplemented with 10 mM DTT and 20% (v/v) glycerol and stored at -80°C until use. This preparation was stable for at least 2 years under these conditions.

### 2.6 In vitro *phosphorylation assays*

Protein phosphorylation was performed by incubating 1 µg of *Ppe*Ald6PRase or myelin binding protein (MBP; Sigma-Aldrich) with the corresponding protein kinase in a standard medium containing 20 mM Tris-HCl pH 7.2, 5 mM MgCl_2_, 0.5 mM CaCl_2_, 2 mM DTT, and 10 µM ATP. Specific media were used to measure SOS2, CDPK and SnRK1 activities. The SOS2 reaction medium contained 50 mM Tris-HCl pH 7.2, 2 mM DTT, 5 mM MgCl_2_, 0.5 mM CaCl_2_, and 100 µM ATP (Gong et al., 2002); the SnRK1 reaction medium contained 20 mM Tris-HCl pH 8.0, 1 mM EDTA, 20 mM MgCl_2_, and 100 µM ATP (Ikeda et al., 1999); and the CDPK reaction media contained 25 mM Tris-HCl pH 7.5, 0.5 mM DTT, 10 mM MgCl_2_, 0.1 mM CaCl_2_, and 50 µM ATP (Zhang et al., 2005). Enzymatic reactions were performed in a final volume of 20 µL with 1 µCi of [^32^P]γ-ATP (Perkin-Elmer). The reactions were started with the addition of the protein kinase (leaf extract, partially purified extract or recombinant protein) and incubated at 25°C. Reactions were stopped by adding SDS-PAGE sample buffer and heating at 95°C for 10 min. The samples were then resolved by SDS-PAGE. To detect the radioactivity incorporated into the protein substrates, gels were stained with Coomassie Blue R-250, dried in a GD-2000 gel dryer system (Hoefer), and exposed to a Storage Phosphor screen (GE Healthcare), following the manufacturer’
ss instructions. Then, the screen was scanned with a Typhoon 9400 (GE Healthcare). Band intensity was quantified using ImageJ (https://www.imagej.nih.gov/ij) and normalized to the intensity of the control band.

We performed a time-course assay to determine lineal reaction conditions using the PPKE with *Ppe*Ald6PRase as substrate. Supplementary Figure S1 shows that lineal conditions were obtained util 20 min of reaction. These conditions were subsequently used for kinetic studies with the PPKE (i.e. determination of *S*_0.5_ and *I*_0.5_ values). Kinetic constants were calculated by plotting PPKE activity versus substrate or effector concentration and fitting experimental data to a modified Hill equation (Ballicora et al., 2007) using Origin 8.0 (OriginLab). *S*_0.5_ and *I*_0.5_ were defined as the concentration of substrate or inhibitor giving 50% of the maximal activity or inhibition, respectively. Data for determining kinetic constants were the mean of at least two independent datasets and were reproducible within a range of ± 10%.

### 2.7 Determination of Ald6PRase activity

Ald6PRase activity was assayed as described by Hartman et al. (2017). Briefly, a proper amount of enzyme was incubated in 100 mM Tris-HCl pH 8.0, 25 mM Glc6P, and 0.25 mM NADPH. Reactions were followed in a final volume of 50 µl at 25°C in a 384-microplate reader (Multiskan GO, Thermo Electron Corporation), following the absorbance at 340 nm. One unit of enzyme activity (U) is defined as the amount of enzyme catalyzing the oxidation of 1 µmol NADPH in 1 min under the specified assay conditions.

### 2.8 Phospho-pendant SDS-PAGE

To analyze the phosphorylation status of Ald6PRase in crude extracts and fractions enriched in phosphoproteins, we employed the Phos-Tag reagent (Wako Chemicals), which delays the migration of phosphorylated proteins in SDS-PAGE. Samples were separated in 10% SDS-PAGE with 100 µM Phos-Tag (following the manufacturer’
ss instructions) or 12% SDS-PAGE without Phos-Tag. After gel washig, proteins were transferred to nitrocellulose membranes at 180 mA for 1h. Membranes were blocked with 5% (w/v) BSA in TBST [50 mM Tris-Cl pH 7.6, 150 mM NaCl, 0.05% (v/v) Tween 20] for 1 h. Primary antibodies against apple (*Malus domestica*) Ald6PRase (Hartman et al., 2017) were diluted 1/1,000 in 1% (w/v) BSA in TBST and membranes were incubated overnight at 4°C. Following extensive washing in TBST, membranes were incubated with HRP-conjugated anti-rabbit IgG (Abcam) diluted 1/10,000 for 1 h. After washing in TBST, membranes were incubated with SuperSignal West Pico Chemiluminescent Substrate (Thermo Fisher Scientific), according to the manufacturer’s instructions. Immunoreactive bands were evidenced in photographic plates (Kodak).

### 2.9 Detection and immunodepletion of protein kinases

SnRK and CDPK in partially purified extracts from peach leaves were evidenced using monoclonal antisera against human phosphorylated-AMPKα (pAMPKα; Cell Signaling Technology, catalog number 2535) and polyclonal antisera against CDPK (Agrisera, catalog number AS132754), respectively.

Protein A-sepharose (Sigma) was washed twice with 5 column volumes of buffer HNTG [20 mM HEPES-NaOH pH 7.5, 150 mM NaCl, 0.1% (v/v) Triton X-100, and 10% (v/v) glycerol]. Sepharose conjugates were aliquoted in 20 µl and antibodies were added (5 µl of polyclonal anti-CDPK and/or 2.5 µl of monoclonal anti-pAMPKα). The resulting samples were incubated for 1 h at RT with gently mixing. Afterward, samples were centrifuged at 3,000 x *g* for 2 min at 4°C. After washing thrice with 1 ml of ice-cold HNTG buffer, the pellets were centrifuged at 3,000 x *g* for 2 min at 4°C. Then, 50 µl of the PPKE were added to each tube and the resulting samples were incubated for 1 h at 4°C with gently mixing. The immunoprecipitated complexes were collected by centrifugation at 3,000 x *g* for 2 min at 4°C, whereas the supernatant was used for the phosphorylation reaction. For sequential immunodepletion, the obtained final supernatant was subjected to the second round of immunodepletion with the second antibody, as described above. A control was processed in parallel without adding antibodies to the mixture and a PPKE control was kept on ice. Results were normalized to the processed control.

### 2.10 Production and purification of recombinant proteins

*Ppe*Ald6PRase was expressed in *E. coli* BL21 Star (DE3) and purified as previously described (Hartman et al., 2017). *Arabidopsis thaliana* KIN10 (*Ath*KIN10; SnRK1, Ca^2+^-independent), *Malus domestica* SOS2 (*Mdo*SOS2; SnRK3, Mg2+-dependent and Ca^2+^-enhanced activity), and *Solanum tuberosum* CDPK1 (*Stu*CDPK1; CDPK, Ca^2+^-dependent) were expressed in *E. coli* BL21 Star (DE3) and purified as previously described (Ferrero et al., 2020; Rojas et al., 2018).

### 2.11 Identification of phosphorylation sites by LC-MS/MS

phosphoproteins were enriched using IMAC-Fe^3+^ chromatography, as described in section 2.4, to identify the putative phosphorylation site on Ald6PRase. The fraction eluted at pH 8.9 was subjected to SDS-PAGE supplemented with Phos-Tag, as described in section 2.8. The gel was stained using colloidal Coomassie Blue G-250 and the band corresponding to the putative phosphorylated Ald6PRase was excised from the gel. In-gel cysteine alkylation and trypsin digestion was performed as previously described (Rosello et al., 2017). Briefly, the gel bands were incubated with 10 mM dithiothreitol (1h at 56 °C) and then with 55 mM iodoacetamide at room temperature for 45 minutes. In-gel proteolytic digestion was performed overnight at 37°C using either sequencing-grade trypsin (Promega) or EndoGluC (Roche). The peptides were extracted at room temperature with a solution containing 60% (v/v) acetonitrile and 0.1% (v/v) trifluoroacetic acid. After a desalting step using C18 microcolumns (ZipTip C18, Millipore), the peptide mixture was vacuum dried and resuspended in 0.1% (v/v) formic acid before injection into a nano-HPLC (UltiMate 3000, Thermo) coupled to a Q-Orbitrap mass spectrometer (Q Exactive Plus, Thermo). Peptide separation was perfomed into a 75 μm × 50 cm, PepMap RSLC C18 analytical column (2 μm particle size, 100 Å, Thermo) at a flow rate of 200 nL/min using a 115 minutes gradient from 1% to 50% (v/v) acetonitrile in 0.1% (v/v) formic acid. Phosphopeptide identification was performed using Proteome Discoverer software (v.1.3.0.339, Thermo) with Sequest as search engine and a database of *Prunus persica* (downloaded from UniProt on September 2018). Precursor mass tolerance was set to 20 ppm, oxidation of Met and phosphorylation of Ser, Thr or Tyr were set as variable modifications while carbamidomethylation of Cys was set as fix modfication. Peptide spectrum matches were filtered to achieve a FDR ≤ 1%. PhosphoRS was used to predict the most probable phosphosite.

## 3. Results

To investigate the putative occurrence of Ald6PRase phosphorylation in peach leaves we performed an acrylamide pendant Phos-Tag SDS-PAGE of crude extracts followed by a western blot. The Phos-Tag reagent reduces the mobility of phosphorylated proteins, which can be reverted if samples are treated with phosphatases. Thus, total proteins from peach leaves were incubated in the absence or presence of CIAP and then separated by SDS-PAGE, with and without Phos-Tag, transferred to a nitrocellulose membrane, and incubated with an anti-Ald6PRase andibody. Figure 1A shows that, in the presence of Phos-Tag, the CIAP-treated sample displayed a major band around 35 kDa corresponding to the non-phosphorylated Ald6PRase (P-). The control sample showed the same band and a shifted band attributable to the phosphorylated Ald6PRase (P+). This additional band was not observed in the SDS-PAGE without Phos-Tag (regardless of the CIAP treatment, Figure 1A).

**Figure 1.**
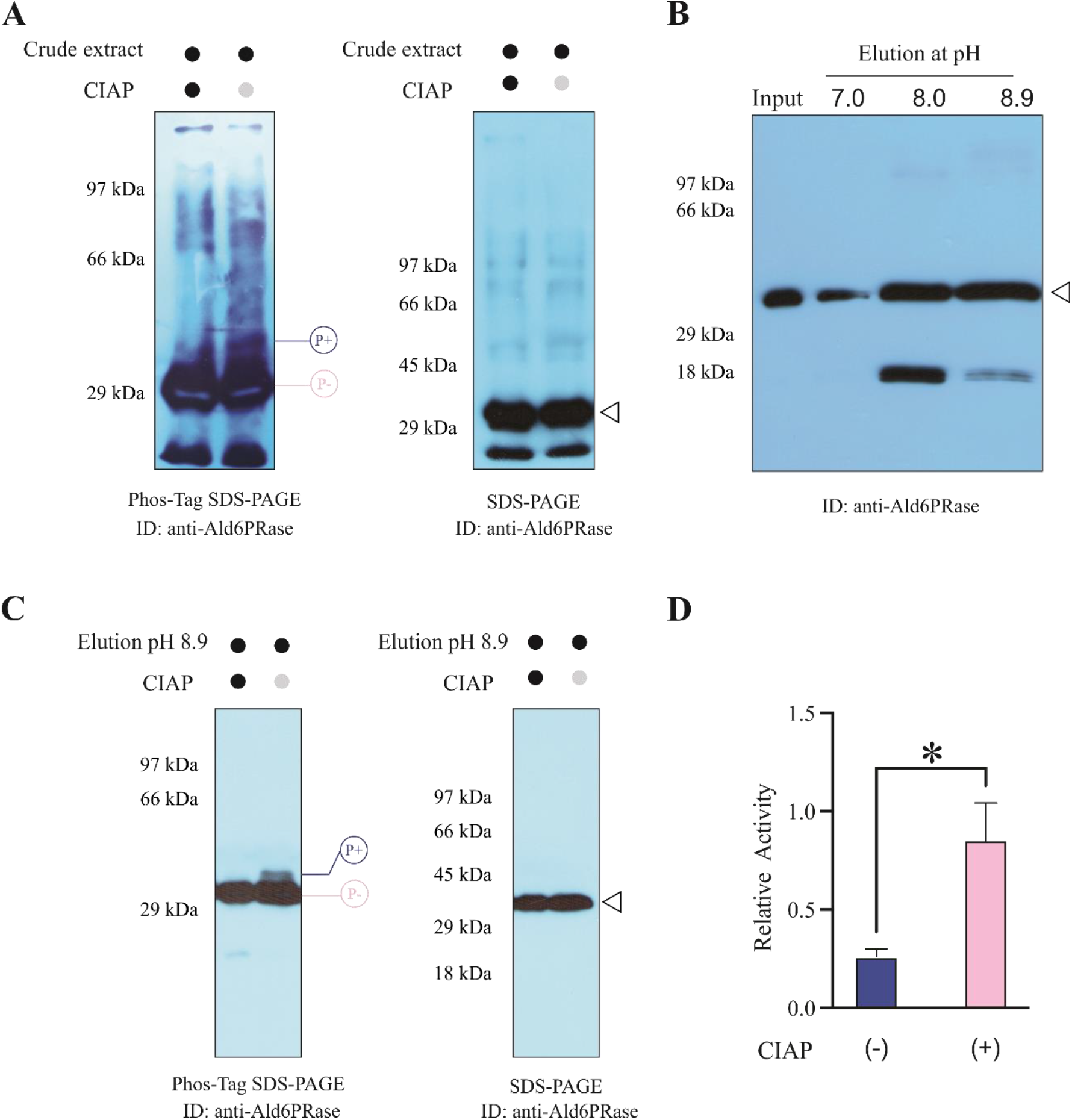
Ald6PRase from peach leaves is phosphorylated *in planta*. (A) Immunodetection of Ald6PRase in total protein extracts from peach leaves treated with CIAP and separated by 12% SDS-PAGE or 10% Phos-Tag SDS-PAGE. (B) Immunoblot of the different elutions obtained from the IMAC-Fe^3+^ chromatography performed with total leaf protein extracts. (C) Immunodetection of Ald6PRase in the pH 8.9 elution from the IMAC-Fe^3+^ chromatography treated with CIAP for 1 h, separated by 12% SDS-PAGE or 10% Phos-Tag SDS-PAGE. (D) Ald6PRase activity of the samples shown in (C).

To further investigate the putative phosphorylation of Ald6PRase *in planta*, we performed an IMAC-Fe^3+^ chromatography, which enriches the sample in phosphorylated proteins (Potel et al., 2018). We loaded a protein extract from peach leaves onto the IMAC-Fe^3+^ column previously equilibrated at pH 6.0. Retained proteins were eluted by increasing the pH (phosphoproteins are expected to elute at alkaline pH values). Ald6PRase was mainly detected in the fractions eluted at pH 8.0 and 8.9, although it was also observed, to a minor extent, in the fraction eluted at pH 7.0 (Figure 1B).

To rule out the possibility of unspecific binding of non-phosphorylated proteins to the IMAC-Fe^3+^ column, the leaf extract was treated with CIAP and then loaded onto the IMAC-Fe^3+^ column. The number of proteins recovered in the phosphoprotein-enriched fraction (pH 8.9) was substantially decreased compared to control samples (Supplementary Figure S2A). Additionally, to discard any indirect interaction of non-phosphorylated Ald6PRase with the IMAC-Fe^3+^ resin (e.g. through a phosphorylated interactor), we performed a denaturing extraction (to abolish protein-protein interactions) followed by IMAC-Fe^3+^ chromatography. In this case, Ald6PRase was observed in the fractions eluted at pH 8.0 and 8.9 (Supplementary Figure S2B), as observed for the native extractions (Figure 1B). These results confirmed that phosphorylated-Ald6PRase directly interacted with the matrix of the IMAC-Fe^3+^ column.

To corroborate that proteins eluted at pH 8.9 from the IMAC-Fe^3+^ column (Figure 1B) were indeed phosphorylated, the fraction was incubated in the absence or presence of CIAP, then analyzed by SDS-PAGE supplemented with Phos-Tag, followed by a western blot. In this experiment, we observed a discrete shift in the mobility of Ald6PRase in the control compared with the CIAP-treated sample that showed only one band (Figure 1C). This result reinforces those obtained with crude extracts (Figure 1A); again, the band-shift was absent regardless of the CIAP treatment in the regular SDS-PAGE (Figure 1C). To test wether phosphorylation affected the enzymatic activity of Ald6PRase, we incubated samples eluted at pH 8.9 from the IMAC-Fe^3+^ column for 1 h in the absence or presence of CIAP. Interestingly, we found a 3-fold increase of Ald6PRase activity in the dephosphorylated samples, suggesting that phosphorylation inhibited Ald6PRase activity (Figure 1D).

To identify the phosphorylated residue/s on Ald6PRase, we scaled up the IMAC-Fe^3+^ chromatography, and the fraction eluted at pH 8.9 was subjected to Phos-Tag SDS-PAGE. The Ald6PRase shifted band was excised from the gel and further analyzed by nano LC-MS/MS. Using EndoGluC as the proteolytic enzyme we could only detect a triply charged phosphorylated peptide (XCorr 4.33 and pRScore 113) that matched the sequence 2-STITLNNGFEMPVIGLGLWRLE-23, located at the N-terminus of peach Ald6PRase (Supplementary Figure S3). The phosphorylation of the N-terminal sequence was further confirmed using digestion with trypsin, as the sequence 2-STITLNNGFEMPVIGLGLWR-21 was detected as a triply charged monophosphorylated ion. However, the information contained in the mass spectra did not allow us to discern which residue (Ser^2^, Thr^3^ or Thr^5^) was phosphorylated.

To gain biochemical information on the protein kinase responsible for Ald6PRase phosphorylation, we partially purified protein kinases from peach leaves, as described in section 2.5. The resulting fraction (PPKE) was kinetically characterized using *Ppe*Ald6PRase as substrate. The *S*_0.5_ values for ATP, Mg^2+^ and Ca^2+^ were 12.7 ± 0.5 μM, 2.6 ± 0.4 mM, and 1.8 ± 0.3 mM, respectively (Supplementary Figure S4A-C). Strikingly, Ald6PRase phosphorylation was maximal in the presence of both Mg^2+^ and Ca^2+^ (Supplementary Figure S4D).

Plant protein kinases (particularly, SnRK1) are regulated by metabolites (Nunes et al., 2013; Piattoni et al., 2011; Toroser et al., 2000; Zhai et al., 2018; Zhang et al., 2009); thus, we tested the effect of several metabolites on the activity of the PPKE. To this end, we used both *Ppe*Ald6PRase and myelin basic protein (MBP; a universal protein kinase substrate) as substrates. Table 1 shows that several metabolites inhibited the activity of the PPKE, irrespective of the protein used as substrate. From these, Glc6P produced ∼25% inhibition; 3-phosphoglycerate (3PGA), Gol6P, PEP and ADP-Glc produced ∼50% inhibition; and pyrophosphate (PPi) produced almost complete inhibition (Table 1). Afterward, some inhibitors were assayed at different concentrations to calculate their *I*_0.5_ values. Table 2 shows that Glc6P displayed the highest *I*_0.5_; PEP, Gol6P, and Fru1,6bisP showed intermediate *I*_0.5_ values; and PPi had the lowest *I*_0.5_.

**Table 1.** Effect of different metabolites on the activity of the PPKE using *Ppe*Ald6PRase and MBP as substrates. All metabolites were tested at 5 mM, except for Tre6P and Fru2,6bisP, whose concentration was 0.1 mM. Data are the mean of two independent experiments ± standard error.

| Metabolite | *PpeAld6PRase*<br>phosphorylation<br>(%) | MBP<br>phosphorylation<br>(%) |
| --- | --- | --- |
| None | 100 | 100 |
| Tre6P | 97 ± 5 | 110 ± 10 |
| Fru2,6bisP | 83 ± 14 | 99 ± 18 |
| Gol6P | 57 ± 11 | 31 ± 4 |
| Glc6P | 74 ± 19 | 79 ± 5 |
| Fru1,6bisP | 28 ± 5 | 38 ± 2 |
| 3PGA | 54 ± 17 | 73 ± 12 |
| PEP | 42 ± 17 | 44 ± 34 |
| ADPGlc | 49 ± 19 | 56 ± 6 |
| PPi | 1.1 ± 0.1 | 3 ± 2 |

**Table 2.** Kinetic parameters for selected metabolites using the PPKE and *Ppe*Ald6PRase as substrate. Kinetic constants were determined using data from two independent experiments.

| Metabolite | $I_{0.5}$ (mM) |
| --- | --- |
| Glc6P | $7.9 \pm 0.3$ |
| PEP | $2.1 \pm 0.1$ |
| Gol6P | $3.3 \pm 0.2$ |
| Fru1,6bisP | $1.9 \pm 0.1$ |
| PPi | $0.49 \pm 0.04$ |

Based on phosphoproteomics data (Supplementary Figure S3) and considering that *in vitro* phosphorylation of recombinant Ald6PRase by the PPKE was maximal in the presence of Ca^2+^ (Supplementary Figure S4D), the following experiments were focused on CDPKs and SnRKs, which belong to the superfamily of Ser/Thr kinases and use Ca^2+^ as a cofactor or are involved in Ca^2+^ signaling, respectively (Hrabak et al., 2003; Wu et al., 2017). Using specific antibodies, we found that both CDPKs and SnRKs were present in the PPKE (Figure 2A). Then, we tested whether these kinases were directly involved in Ald6PRase phosphorylation. As shown in Figure 2B, immunodepletion of both kinases reduced *Ppe*Ald6PRase phosphorylation by approximately 50% compared with the untreated control, thus confirming that both kinase types played a role in Ald6PRase phosphorylation.

**Figure 2.**
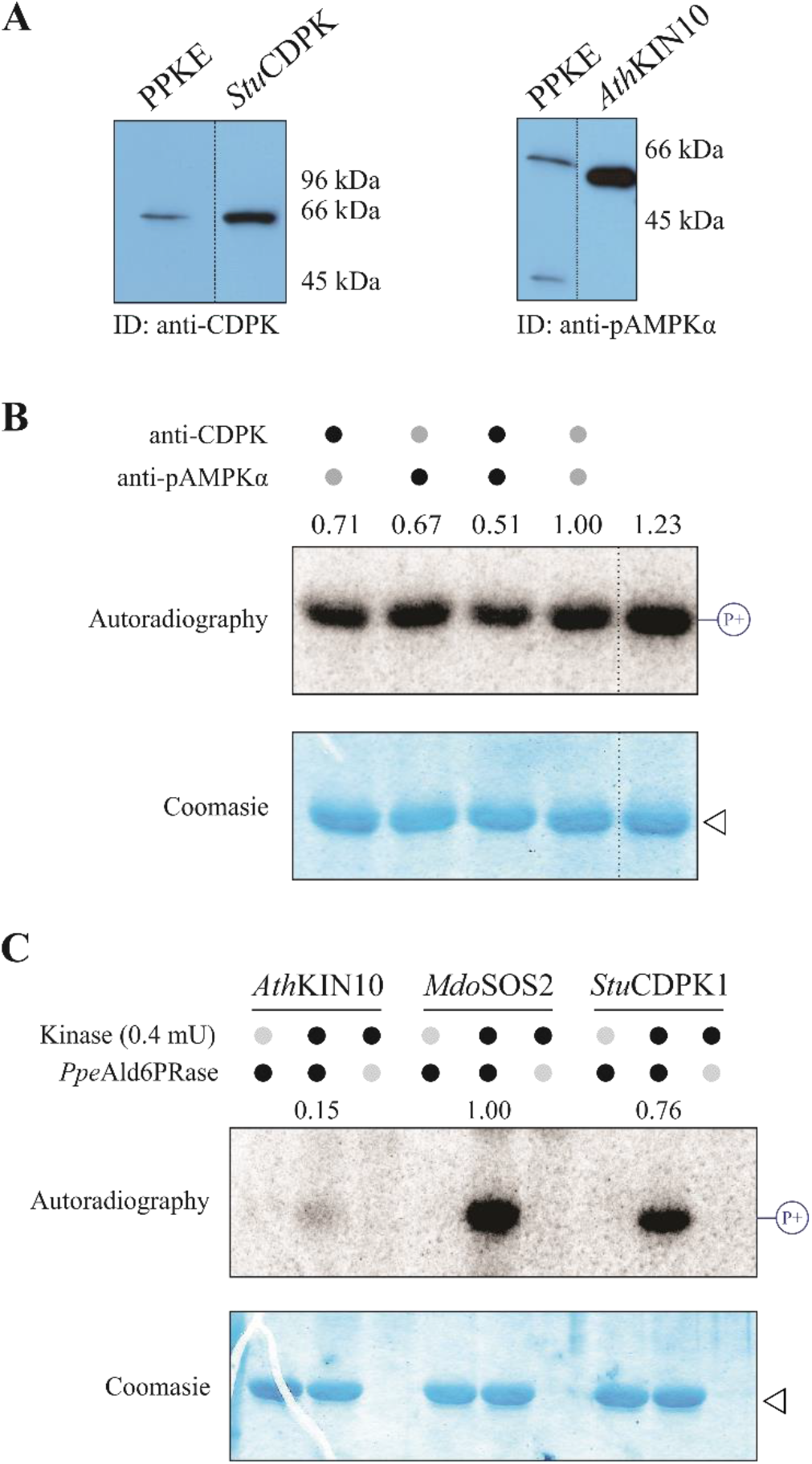
The PPKE contains protein kinases from the CDPK and SnRK families. (A) Immunodetection of SnRK and CDPK kinases within the PPKE. (B) Immunodepletion of CDPK (lane 1), SnRK (lane 2), and consecutive CDPK and SnRK (lane 3) from the PPKE and further *in vitro* phosphorylation of *Ppe*Ald6PRase. The controls were PPKE treated with no antibody (lane 4) and PPKE kept on ice during the whole procedure and used immediately before the phosphorylation reaction (lane 5). (C) *In vitro* phosphorylation of *Ppe*Ald6PRase using 0.4 mU of *Ath*KIN10, *Mdo*SOS2 and *Stu*CDPK1.

To further investigate the specificity of SnRKs and CDPKs to phosphorylate Ald6PRase, we reconstituted three different systems using, alternatively, recombinant *Ath*KIN10, *Mdo*SOS2, and *Stu*CDPK1 (Vlad et al., 2008). Phosphorylation assays were performed *in vitro* using a fixed amount of protein kinase activity (0.4 mU) to phosphorylate *Ppe*Ald6PRase. Phosphorylation of *Ppe*Ald6PRase was maximal with *Mdo*SOS2, slightly lower with *Stu*CDPK1 and almost negligible with *Ath*KIN10 (Figure 2C). Altogether, our results strongly suggest that peach Ald6PRase is a substrate of both SOS2 and CDPK.

## 4. Discussion

Sugar-alcohols are primary photosynthetic products in plants from the Rosaceae family (Figueroa et al., 2016; Loescher and Everard, 2004). Therefore, the first enzymatic step diverting Glc6P into sugar-alcohol metabolism must be a critical control point. In other words, Ald6PRase should be tightly regulated to balance carbon flux between different metabolic pathways, including sucrose and starch synthesis, respiration and demands from heterotrophic tissues (Teo et al., 2006). Transcription of the gene encoding Ald6PRase is regulated by environmental and physiological cues, such as cold and ABA (Kanayama et al., 2006; Lloret et al., 2017; Lou et al., 2018; Suzuki and Dandekar, 2014). Additionally, the enzymatic acitiviy of Ald6PRase is inhibited by hexose-phosphates (Glc1P, Fru1,6bisP and Fru6P), inorganic orthophosphate and oxidizing agents (Hartman et al., 2017). The latter poses AldPRase as a key player in carbon channelling to the synthesis of sugar-alcohols under stressful conditions, when preservation of NADPH (which provides the reducing equivalents in the reaction catalyzed by Ald6PRase) is critical for maintaining antioxidant systems in their reduced (and active) form (Del Corso et al., 1994). Thus, Ald6PRase has to balance NADPH consumption with the production of the sugar-alcohol, which functions as osmoprotectant and radical scavenger (Figueroa et al., 2016; Loescher and Everard, 2004).

This work demonstrates that Ald6PRase from peach leaves is phosphoryated *in planta*. In crude extracts from illuminated leaves, the phosphorylated form accounted for a small fraction of the total Ald6PRase pool. It has been suggested that low-stoichiometric PTMs might reflect the occurrence of time- and/or place-specific modifications (Prus et al., 2019). Thus, if Ald6PRase is differentially regulated in particular cell types, the phosphorylated pool would represent a minor fraction of the protein in the whole leaf extract. Besides this, it is important to note that the activity of the CIAP-dephosphorylated Ald6PRase was considerably (3-fold) higher than the phosphorylated form. This means that phosphorylation of the enzyme could be considered an important regulatory mechanism to modulate the synthesis of glucitol in leaves.

Our mass spectrometry analysis showed that phosphorylation of peach Ald6PRase occured at the N-terminus. Currently, little is known regarding the phosphorylation of proteins from the AKR superfamily. Rat AKR1A1 is phosphorylated at Ser^4^ (Lundby et al., 2012), which is strikingly similar to the N-terminal phosphorylation found on peach Ald6PRase. In different mammalian cell lines, the agonist-mediated stimulation of protein kinase C activity increases the amount of the phosphorylated form of aldose reductase and favors a switch from the cytosol to mitochondria (Varma et al., 2003). Direct phosphorylation of recombinant aldose reductase by protein kinase C was demonstrated *in vitro*, although no experiments were performed to evaluate neither the effect of this PTM on the kinetic properties of the enzyme nor the identity of the phosphorylated residue explored (Varma et al., 2003). The PhosPhat database (Durek et al., 2009; Heazlewood et al., 2008; Zulawski et al., 2013) indicates that the Arabidopsis At2G21260 protein, an ortholog of peach Ald6PRase with demonstrated Ald6PRase activity (Rojas et al., 2019), is phosphorylated at Thr^256^. Our mass spectrometry analysis failed to identify whether or not the corresponding residue in peach Ald6PRase (Thr^257^) was phosphorylated. A plausible explanation for these differences is that plant AKRs could be subject to multiple phosphorylation events, depending on the cell type and/or the metabolic scenario, as it has been reported for other enzymes involved in carbon metabolism (Chollet et al., 1996; Huber and Huber, 1996; Nimmo, 2003). It is worth mentioning that phosphorylated At2G21260 was obtained from Arabidopsis cell cultures grown under nitrogen starvation (Durek et al., 2009; Heazlewood et al., 2008; Zulawski et al., 2013), whereas *Ppe*Ald6PRase was obtained from mature, illuminated peach leaves, which could easily account for the distinct phosphorylation patterns observed for these two orthologs.

Using a PPKE we found that phosphorylation of *Ppe*Ald6PRase is enhanced in the presence of Ca^2+^. Immunodetection assays showed that the PPKE contained protein kinases from the SnRK and CDPK families. Furthermore, immunodepletion of both kinase types from the PPKE greatly reduced Ald6PRase phosphorylation, indicating that these proteins were, at least in part, responsible for Ald6PRase phosphorylation. However, the kinase activity remaining in the PPKE after immunodepletion of SnRKs and CDPKs points to a third type of kinase capable of phosphorylating Ald6PRase, the nature of which remains to be elucidated.

The allosteric regulation of plant SnRKs has been well described; for instance, SnRK1 from Arabidopsis is inhibited by Glc6P, Glc1P, and trehalose 6-phosphate (Nunes et al., 2013; Piattoni et al., 2011; Toroser et al., 2000). However, to the best of our knowledge, no reports show that CDPKs are allosterically regulated. Interestingly, the kinase activity of the PPKE was inhibited by several metabolites, namely Glc6P, PEP, Gol6P, Fru1,6bisP and PPi. Notably, both the substrate (Glc6P) and the product (Gol6P) of the reaction catalyzed by Ald6PRase inhibited the kinases present in the PPKE with *I*_0.5_ values within the low millimolar range. The cytosolic concentration of Glc6P in peach leaves has been estimated to be 9.5 mM (Hartman et al., 2017), a value close to the *I*_0.5_ obtained for Glc6P with the PPKE. We could not find experimental data to estimate the concentration of Gol6P in peach leaves; however, the amount of Glc6P in apple fruits is almost 10-fold higher than Gol6P (Zhang et al., 2017). Thus, we speculate that Glc6P (but not Gol6P) could be considered a physiologically relevant inhibitor of the protein kinases in peach leaves.

The phosphorylation of Ald6PRase by kinases from the CDPK and SnRK families was further confirmed using an *in vitro* system. Many proteins have been described as CDPK targets (Curran et al., 2011); however, none of them are members of the AKR superfamily. Wei et al. (2016) identified 123 CDPK genes from five rosaceous species (apple, pear, peach, strawberry and plum), demonstrating the complexity of the CDPK family in these plant species. CDPKs have been described as positive regulators of abiotic stress responses; moreover, over-expression of CDPKs was associated with enhanced stress tolerance (Asano et al., 2012; Boudsocq and Sheen, 2013). The modulation of Ald6PRase activity by CDPKs could be related to stress tolerance events, although more evidence is needed to support this hypothesis.

Besides the plasma membrane Na^+^/H^+^ antiporter SOS1 (Qiu et al., 2002), only one putative target of SOS2 (a member of the SnRK3 clade) has been described so far, the ADP-glucose pyrophosphorylase from wheat endosperm (Ferrero et al., 2020). In this work, we found a second putative target of SOS2 (i.e. peach Ald6PRase), which, as in the case of ADP-glucose pyrophosphorylase, is involved in carbon metabolism. The SOS pathway plays a crucial role in the response to salt stress (Qiu et al., 2002); interestingly, Gol is also involved in salt stress tolerance (Pommerrenig et al., 2007). Recently, Yu et al. (2021) described the phosphorylation and subsequent activation of peach fruit GolDHase by SnRK1. Based on these findings, we postulate that SnRKs are significant regulators of Gol metabolism *in vivo*.

Overall, our results shed light on the putative regulation of Ald6PRase by phosphorylation in illuminated mature peach leaves (Figure 3). Such modification seems to occur at the N-terminus and would be performed by a CDPK and/or SnRK. Dephosphorylation increased the enzymatic activity of Ald6PRase obtained from mature, illuminated leaves; in other words, phosphorylation inhibits Ald6PRase activity (Figure 3). In addition, the PPKE was inhibited by hexose-phosphates, whose levels vary along the diel cycle, being higher in the light than in the dark (Arrivault et al., 2009). Considering this metabolic scenario, we speculate that these kinases are inhibited during the day and active at night. This assumption implies that phosphorylation of Ald6PRase is minimal in the light and maximal in the dark. Consequently, the activity of Ald6PRase would be higher during the day than at night (Figure 3). Indeed, the levels of Gol in leaves fluctuate during the diel cycle, with a maximum at the end of the day, followed by a marked decrease after dusk (Chong and Taper, 1971; Li et al., 2007). Then, it seems feasible that phosphorylation plays a major role in controlling the levels of leaf Gol. Furthermore, phosphorylation of other enzymes, such as sucrose-6-phosphate synthase (Huber and Huber, 1996; Toroser et al., 2000), would orchestate primary carbon metabolism by balancing the levels of sucrose and Gol produced during the diel cycle.

**Figure 3.**
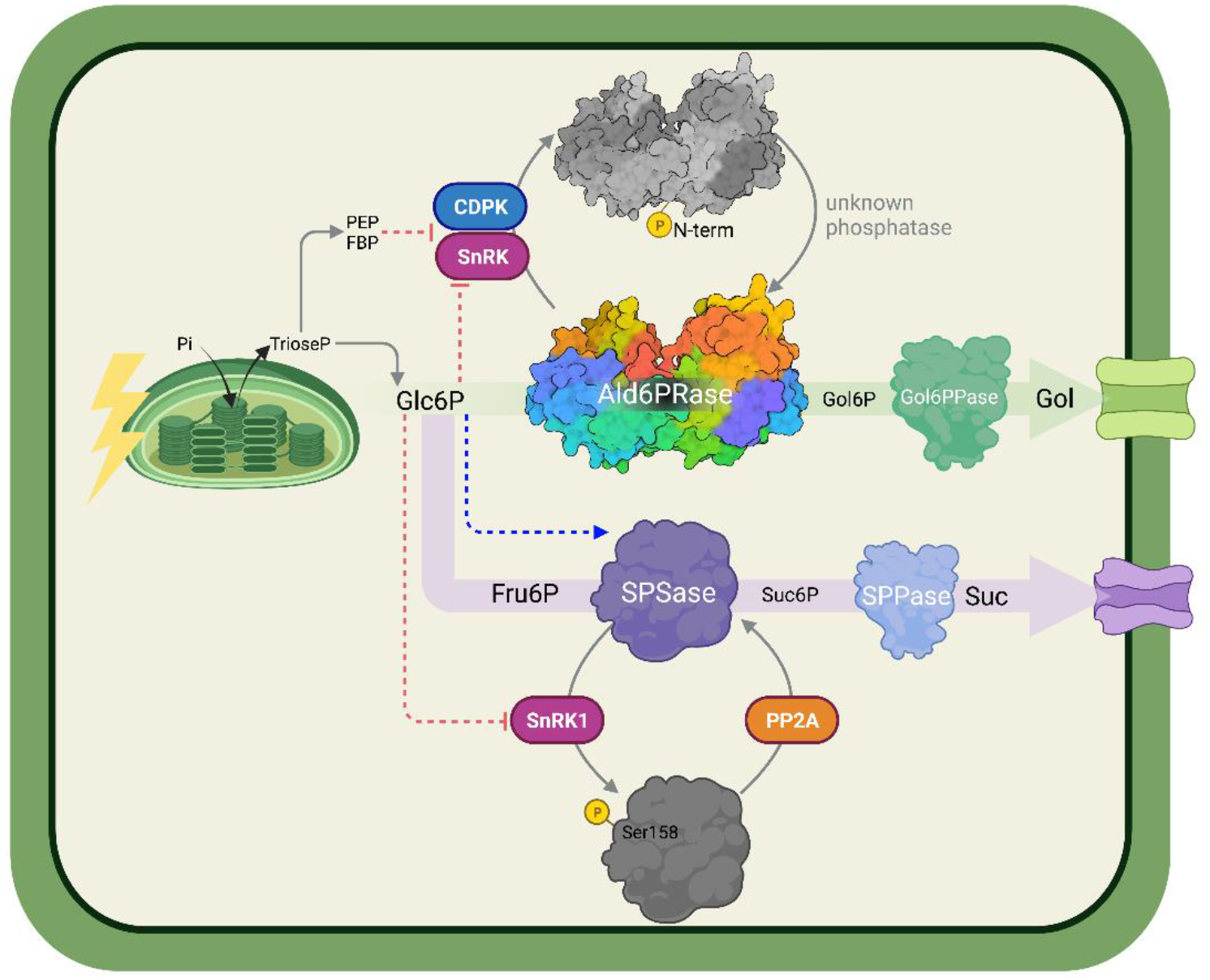
Proposed model for the phosphorylation-dependent regulation of primary carbon metabolism in peach leaves. The activities of Ald6PRase and sucrose-phosphate synthase (SPSase) are inhibited by phosphorylation. The activity of Ald6PRase could be restored by a yet unknown kinase, while a PP2A phosphatase dephosphorylates SPSase. In both cases, kinases are inhibited by Glc6P, namely the substrate of Ald6PRase, and the precursor of substrates used by SPSase. High levels of Glc6P (and other high-energy metabolites, such as PEP and Fru1,6bisP) would relieve Ald6PRase and SPSase activity by inhibiting protein kinases, thus allowing the synthesis of Gol and sucrose, respectively. Red dotted lines, inhibition; blue dotted lines, activation.

## 5. Conclusion

In this work, we showed that Ald6PRase is phosphorylated in planta. Additionally, we found that sugar-phosphates inhibit protein kinases from peach leaves. We also demonstrated that *Ppe*Ald6PRase was phosphorylated *in vitro* by recombinant CDPKs (*Md*SOS2 and *Stu*CDPK1). Overall, our results strongly suggest that phosphorylation is an important mechanism for regulating the activity of Ald6PRase. These findings contribute to a better understanding of the fine-tuning of enzymes involved in carbon metabolism in peach leaves. Overall, our results could be helpful to engineer mutant enzymes (insensitive to phosphorylation) to manipulate Gol synthesis in other economically important plant species.

## Acknowledgements

MDH is a fellow from Agencia I+D+i. DMLF was a PhD fellow from CONICET. BER is a postdoctoral fellow from CONICET. AAI and CMF are researchers from CONICET. The authors thank Dr. K. Allmeroth for care reading of the manuscript and valuable input.

## Author contributions

Conceptualization: MDH, CMF and AAI; Investigation: MDH, BER, DMLF, CMF, AL, RD. Formal analysis: all authors; Writing - Original Draft: MDH, BER and CMF; Writing - Review & Editing: all authors.

## Abbreviations

Ald6PRase: aldose-6-phosphate reductase
CDPK: calcium-dependent protein kinase
Glc6P: glucose 6-phosphate
Gol: glucitol
Gol6P: glucitol 6-phosphate
SnRK: SNF1-related kinase
SOS2: salt overly sensitive 2

---

**Figure S1.**
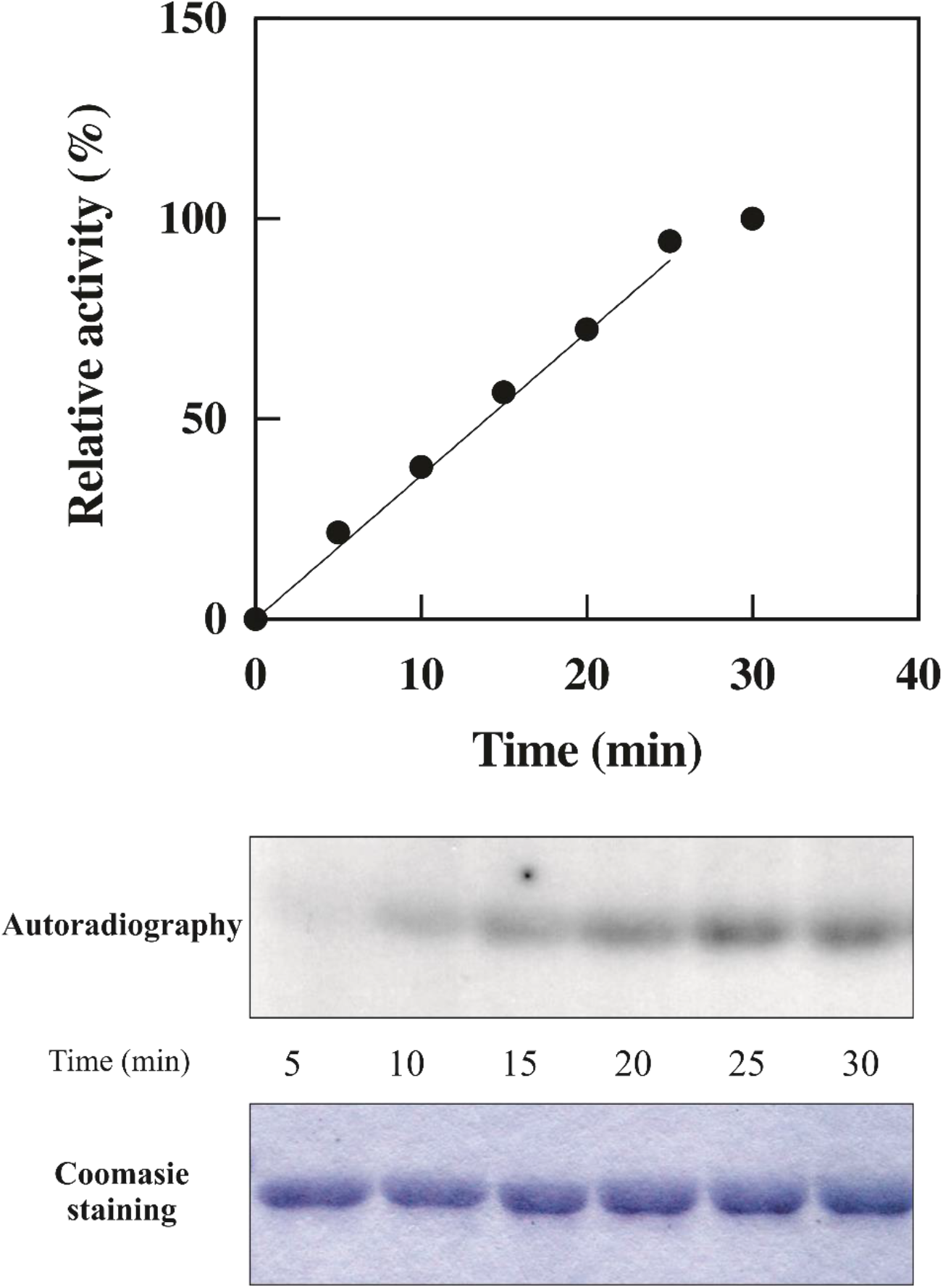
Phosphorylation of Ald6PRase by PPKE is linear up to 20 min. Quantification of [^32^P]γ-ATP incorporated into Ald6PRase in a 30 min time-course.

**Figure S2.**
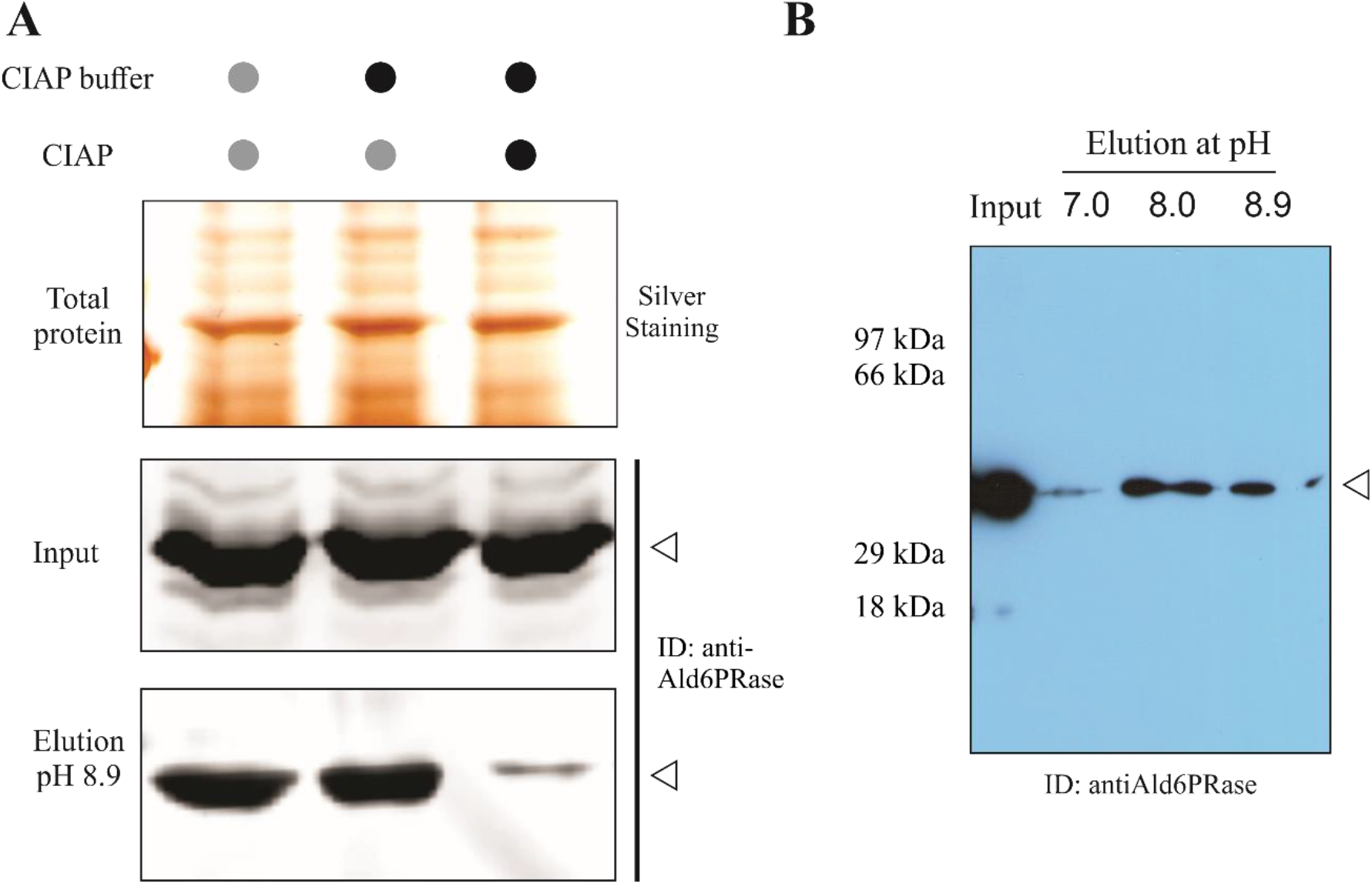
Assays to confirm the phosphorylation status of Ald6PRase *in planta*. (A) The crude extract was dephosphorylated before the IMAC-Fe^3+^ chromatography. Lane 1, protein extract incubated 5 h at 30°C (control); lane 2, protein extract incubated 5 h at 30°C in the presence of CIAP buffer (control); lane 3, protein extract set 5 h at 30°C in the presence of CIAP buffer and 5 U CIAP. Upper panel, silver staining; middle panel, input; bottom panel, elutions at pH 8.9 from the IMAC-Fe^3+^ chromatography. (B) Immunodetection of Ald6PRase in the elutions of the IMAC-Fe^3+^ chromatography performed with a protein extract obtained under denaturing conditions.

**Figure S3.**
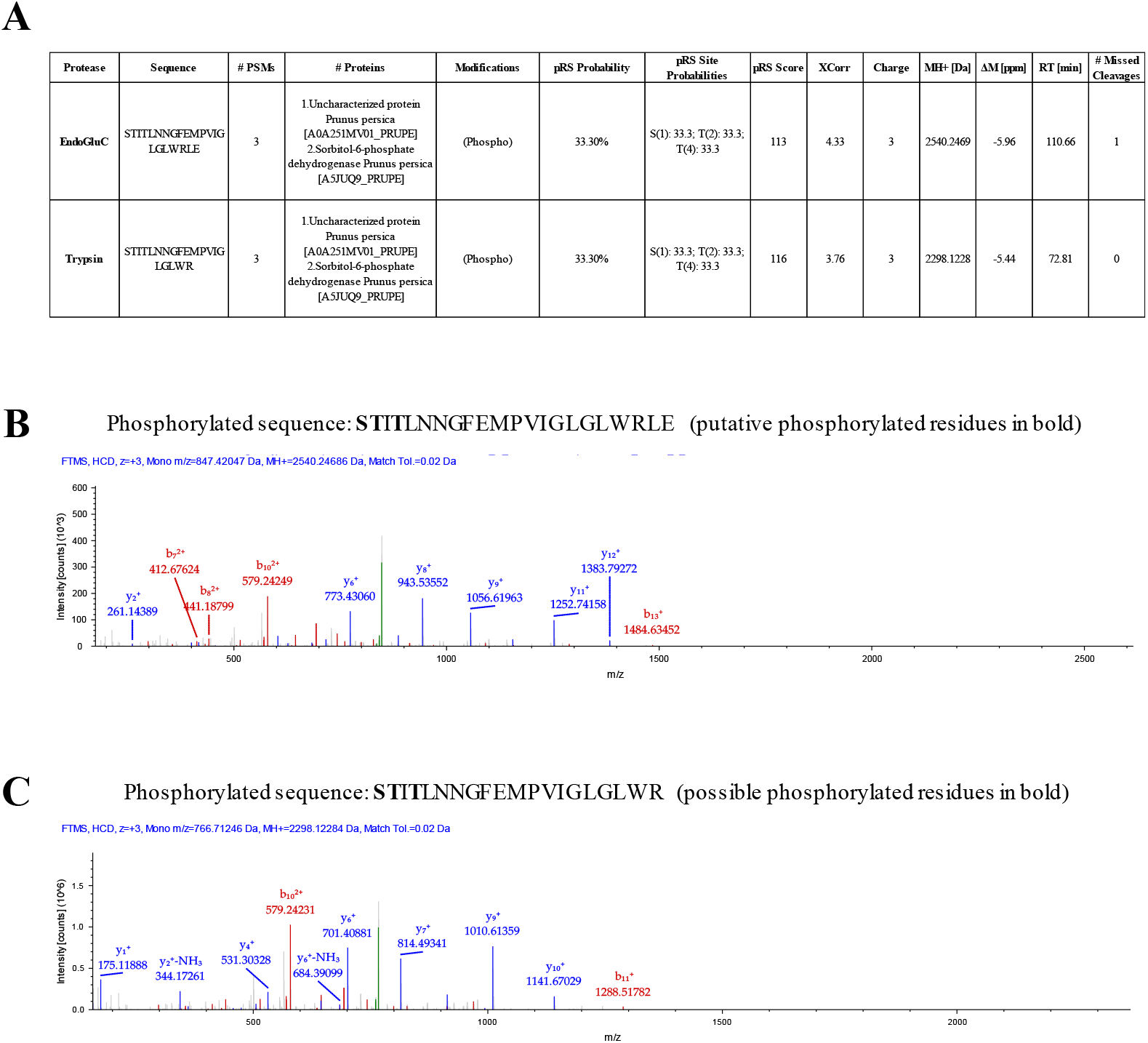
Analysis of the phosphorylation site in Ald6PRase from peach leaves. (A) Information of the phosphorylated peptides obtained from Ald6PRase from peach leaves treated with EndoGluC and Trypsin. (B) MS/MS spectrum of the phosphorylated peptide identified using EndoGluC. (C) MS/MS spectrum of the phosphorylated peptide obtained using Trypsin. Red, b series; blue; y series.

**Figure S4.**
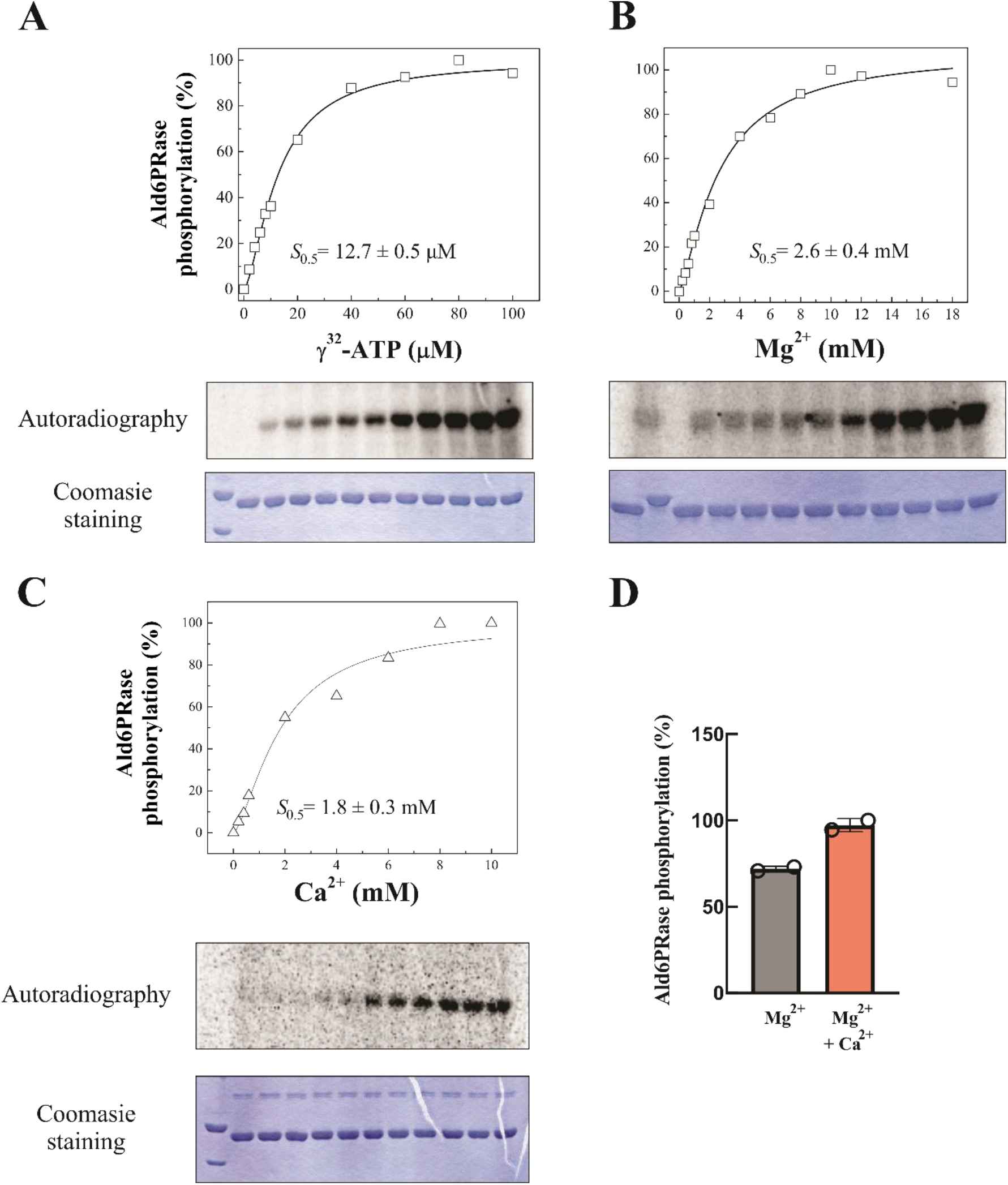
Kinetic characterization of the PPKE. Saturation curves for ATP (A), Mg^2+^ (B), and Ca^2+^ (C). Kinetic constants were calculated as described in section 2.6. (D) Phosphorylation of *Ppe*Ald6PRase by the PPKE in the absence or presence of Ca^2+^.

